# New distance measure for comparing protein using cellular automata image

**DOI:** 10.1101/2023.06.16.545334

**Authors:** Luryane F. Souza, Hernane B. de B. Pereira, Tarcisio M. da Rocha Filho, Bruna A. S. Machado, Marcelo A. Moret

**Affiliations:** Centro de Ciências Exatas e das Tecnologias, Universidade Federal do Oeste da Bahia, Barreiras, Bahia, Brazil; Programa de Modelagem Computacional e Tecnologia Industrial, SENAI-CIMATEC, Salvador, Bahia, Brazil; DEDC, UNEB, Salvador, Bahia, Brazil; Instituto de Física, Universidade de Brasília, Brasília, Distro Federal, Brazil; DCET, UNEB, Salvador, Bahia, Brazil

## Abstract

One of the first steps in protein sequence analysis is comparing sequences to look for similarities. We propose an information theoretical distance to compare cellular automata representing protein sequences, and determine similarities. Our approach relies in a stationary Hamming distance for the evolution of the automata according to a properly chosen rule, and to build a pairwise similarity matrix and determine common ancestors among different species in a simpler and less computationally demanding computer codes when compared to other methods.

## Introduction

Bioinformatics has been growing recently and consolidating itself as a research area, bringing together researchers from different areas such as molecular biology, physics, mathematics, and computer science, among others. Its origins date back to the first protein sequencing studies in the 1950s when insulin was sequenced well before the first microcomputers [1–3]. More recently, there has been a significant increase in sequencing tools, with sequence analysis software such as ClustalW [4] and protein databases as UniProt [5], GenBank [6], PDB [7], and PDB2 [8]. Alongside bioinformatics, molecular biology has grown in recent years with the emergence of different computational approaches designed to study protein folding and sorting [9–11]. A protein is a macromolecule made up of 20 types of amino acids that can represented in its primary form as a string of characters for each amino acid. Different approaches for graphical representations of protein sequences were proposed in Refs. [11–21]. The approach we use in the present work is based on cellular automata (CA) images generated using the sequence of a given protein as initial state. Amino acids are encoded into valid entries of a cellular automaton (see for instance Xiao et al. [17]), using a digital code based on the rules of similarity, complementarity, molecular recognition theory and information theory [16]. A coding based on hydrophobicity indices of amino acids was used in a reduced form by Kavianpour and Vasighi [11] to encode protein sequences to extract features from the images and determine the structural class of the protein. Chaudhuri et al. [20] used a coding with an eight-digit binary code based on the analysis of the molecular structure of each amino acid.

Similarities among sequences, i. e. homologous sequences, hold important information, such as similar functions or as an indication of a recent common ancestor [22]. Indeed, evolutionary relationships between protein sequences can be determined from sequence comparison methods [23–28]. Rahman et al. [21] proposed a method for decomposing CA images using wavelet decomposition and used the horizontal image of this decomposition for protein comparison from an image quality metric. The Hamming distance have been successfully used to evaluate the stability of the concentration of soot during controlled combustion of acetylene and natural gas, within the spatiotemporal standards generated by the evolution of the CA-based system [29]. In previous work by some of the authors [28] we used cellular automata imaging of the Spike proteins to compare variants of the SARS-CoV-2 virus using the stationary Hamming distance, and determined the variants that shared recent common ancestors by looking only at the evolution of the distance between the variant CA and the one for the reference protein initially found in Wuhan-China. As a continuation, we propose a method to build the distance matrix between species pairs using the stationary Hamming distance measuring the dissimilarity between different species. We apply this method for different proteins: ND5, ND6, transferrin, and beta-globin. The proposed approach is effective for grouping similar species and, using cophenetic correlation coefficients, building dendrograms similar to those obtained using the p-distance from the package MEGA [30].

Considering that the p-distance measures the differences between two sequences and that information loss may occur when transforming a protein sequence into a cellular automaton, our results confirm that this loss is minimal, and that our methodology can be used in the analysis of similar proteins. Another advantage is that we use a simple comparison metric not requiring more elaborate processing methods or image textures.

## Materials and methods

A Cellular automaton (CA) is a discrete dynamical system in both space and time evolving under a given spatially local rule. Despite their simplicity they often model complex systems [31]. It is defined from five components: *L, S, N, f* and *B*, with *L* a n-dimensional spatial lattice of cells with values 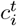, *i* = 1, 2, 3, …, *M* at time *t*. Each cell assumes values in the set of possible states *S*. The neighborhood *N* of a given cell *i* is the set of cells considered in the transition rule *f*. Finally, the boundary conditions of the automaton is represented by *B*.

Here we consider in the presen work one-dimensional cellular automata with a neighborhood of cell *i* it given by the cells *i −* 1, *i* and *i* + 1:

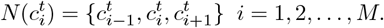

and a set of two possible states *S* = {0, 1}. Therefore, we have a total of 2^3^ = 8 different possible neighborhoods. The transition function *f* expresses the state assumed by each cell in the next time step according to its neighborhood as a query list for each possible neighborhood state. We thus have a total of 2^8^ = 256 possible evolution rules for the cellular automaton, each rule enumerated 0 to 255 given by the decimal form of it binary representation, as exemplified in Fig 1. The boundary condition *B* determining the neighborhood of cells at the extremities of the cellular automaton can be of four types: fixed contour, random, periodic and reflecting [32]. We consider here periodic boundary conditions such that that *c*_*M*+1_ = *c*_1_ and *c*_0_ = *c*_*M*_. Fig 2 illustrates the procedure of forming an image of the cellular automaton composed by the lines for each discrete time value *t* according to the evolution rule.

**Fig 1.**
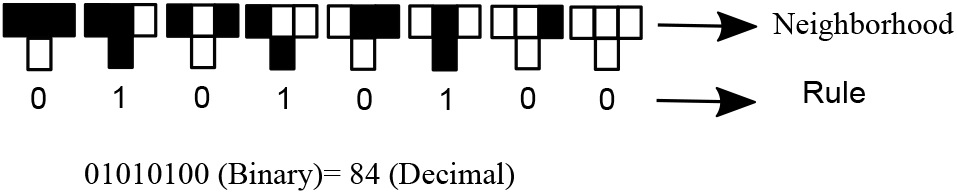
Cellular automaton rule *n*^*o*^ 84 with for the eight types of neighborhoods in the cellular automaton.

**Fig 2.**
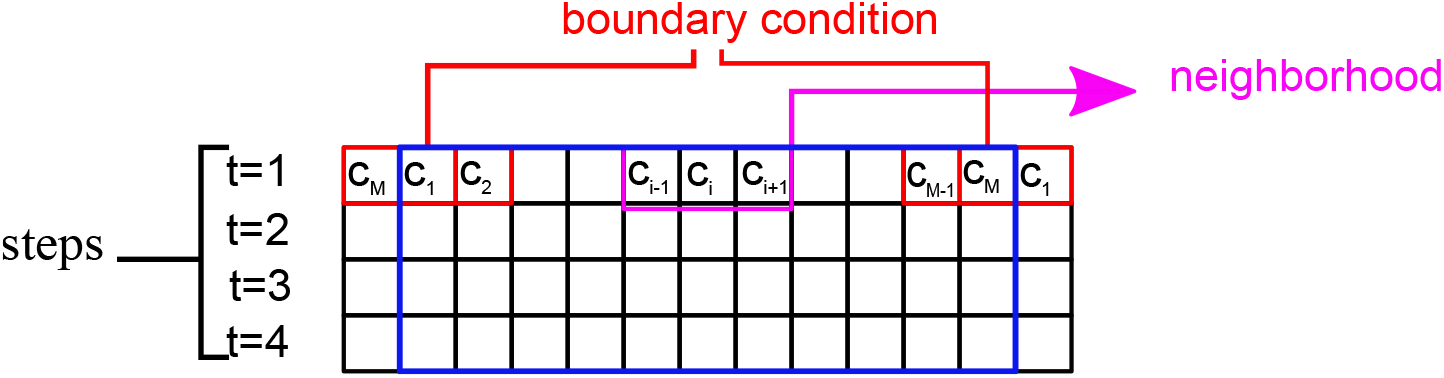
Cellular automaton image

Since we use a two state cellular automaton, each amino acid will be encoded in a binary code, with different possibilities studied in the literature [11, 16, 20, 33]. For the present study we use the code proposed by Chaudhuri et al. [20] and shown in Table 1, which encodes each amino acid with an 8-digit code based on the molecular structures of amino acids. Among other possibilities, this choice is justified by the fact that it yields a better grouping of close species.

**Table 1.** Encoding of amino acids, deletions and missing protein sequence data after alignment.

| Aminoacid | Code | Aminoacid | Code |
| --- | --- | --- | --- |
| Glycine (G) | 00000000 | Cysteine (C) | 01000100 |
| Alanine (A) | 00000100 | Threonine (T) | 00110100 |
| Proline (P) | 00100110 | Asparagine (N) | 00101110 |
| Valine (V) | 00010110 | Glutamine (Q) | 00101111 |
| Methionine (M) | 00110110 | Tyrosine (Y) | 10100100 |
| Tryptophan (W) | 10110110 | Histidine (H) | 01111110 |
| Phenylalanine (F) | 10000100 | Lysine (K) | 00110111 |
| Isoleucine (I) | 00011110 | Arginine (R) | 01111111 |
| Leucine (L) | 00010111 | Aspartic Acid (D) | 01110100 |
| Serine (S) | 00100100 | Glutamic Acid (E) | 01110110 |
| Deletion (-) | 11111111 | Missing information (?) | 11111110 |

The first step is to align the different protein sequences considered such that each associated automata are of the same size. Deletions and losses are identified and properly represented in each automaton by the respective codes in Table 1. Considering a protein of size *P*, the initial condition will have a size of *M* = 8 *× P*. As a first illustration of our approach we show in Fig. 3 the cellular automaton image of Beta-Globin protein for six different animal species, for a total evolution of *t* = 500 steps (this value will be used for the remaining of the present paper).

**Fig 3.**
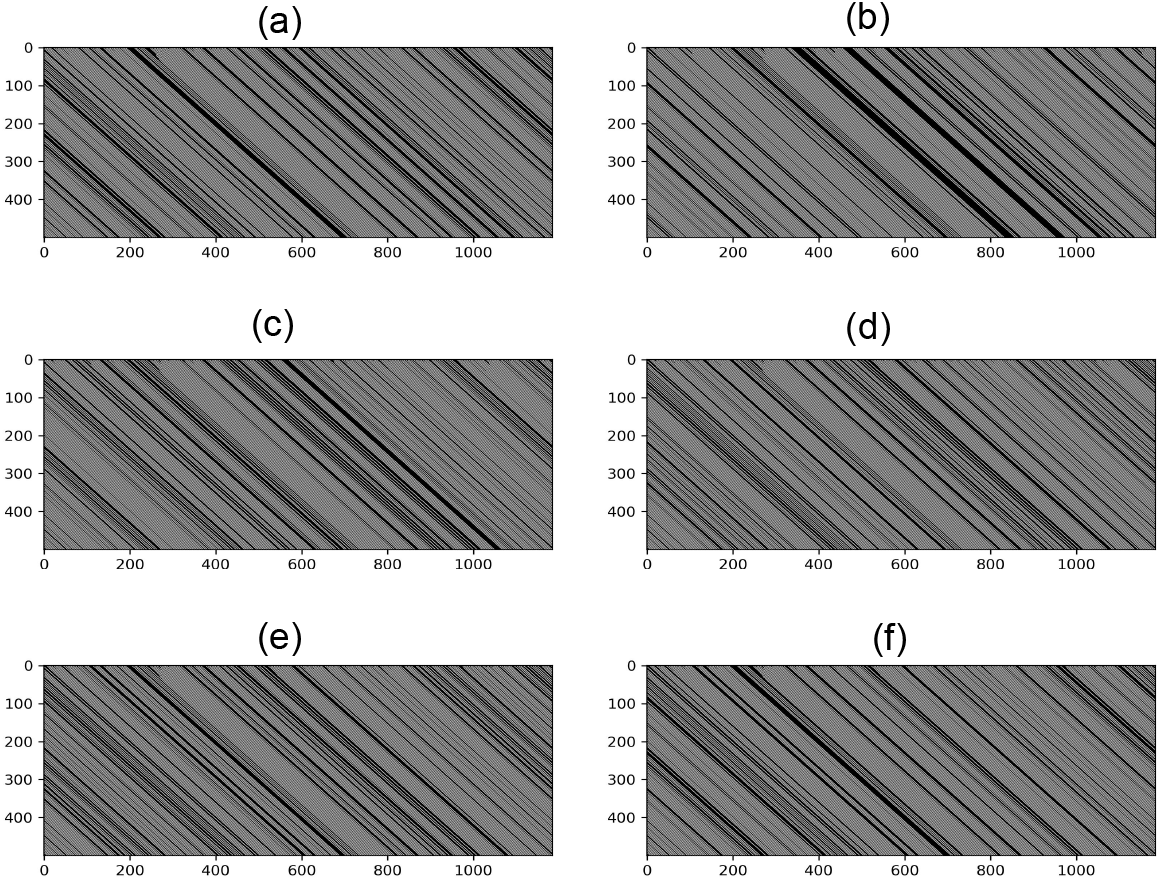
Cellular automaton image of the Beta-Globin protein for A) Human, B) Shark, C) Catfish, D) Turtle, E) Swift and F) Coyote species.

The images of the associated cellular automata provide signatures for different proteins and are used to determine similarities/differences between species. A comparison between those images can be performed with a low computational cost using the Hamming distance from information theory [34] given by the number of changes needed to transform one sequence into another, and implement for the generated images of two cellular automata *CA*_*A*_ and *CA*_*B*_ as:

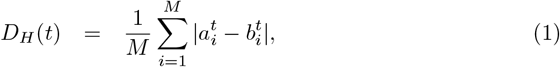

with 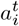 and 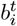 the values of the *i*th cells of automata *CA*_*A*_ and *CA*_*B*_ at step *t*, respectively. The size of the automaton is *M* = 8 *× P*, where *P* is the size of the aligned protein sequence. As shown in the next section the Hamming distance saturates after a relatively small number of steps. We denominate this saturated value the Stationary Hamming Distance (SHD), which is then used to build the similarity matrix.

## Results and Discussion

We apply our approach to the following four protein sequences: beta-globin, NADH Dehydrogenase 5 (ND 5), NADH Dehydrogenase 6 (ND 6) and transferrin. This choice was motivated by the requirement to be able to perform comparisons with previous results in the literature [13, 35]. All sequences were aligned using the ClustalW system [30, 36]. Our results are then compared to those obtained from pairwise p-distance from ClustalW.

### NADH Dehydrogenase 5 (ND 5)

Protein ND 5 is a sub-unit of the mitochondrial respiratory enzyme complex I (NADH: ubiquinone oxidoreductase) [37], and is responsible for mitochondrial electron transport. Mutations and defects in this enzyme can cause Leigh’s disease and MELAS syndrome. Being highly conserved in eukaryotes, we use data from these sequences to analyze similarities between mammalian species. We consider here the following nine species: Human, Gorilla, Pigmy Chimpanzee, Common Chimpanzee, Fin Whale, Blue Whale, Rat, Mouse, and Opossum. All sequences were taken from the NCBI protein database [6], and their identifications are given in S1 Table. The aligned sequences have 613 entries each, and are represented using the coding in Table 1, resulting into a binary sequence of length 4904 for the initial condition of the cellular automaton. The cellular automata image is the generated from the prescription in the previous section and the Hamming distance in Eq. (1) between two species as a function of the number of steps. The results for the distance between each of the nine species and Humans are shown in Fig. 4.

**Fig 4.**
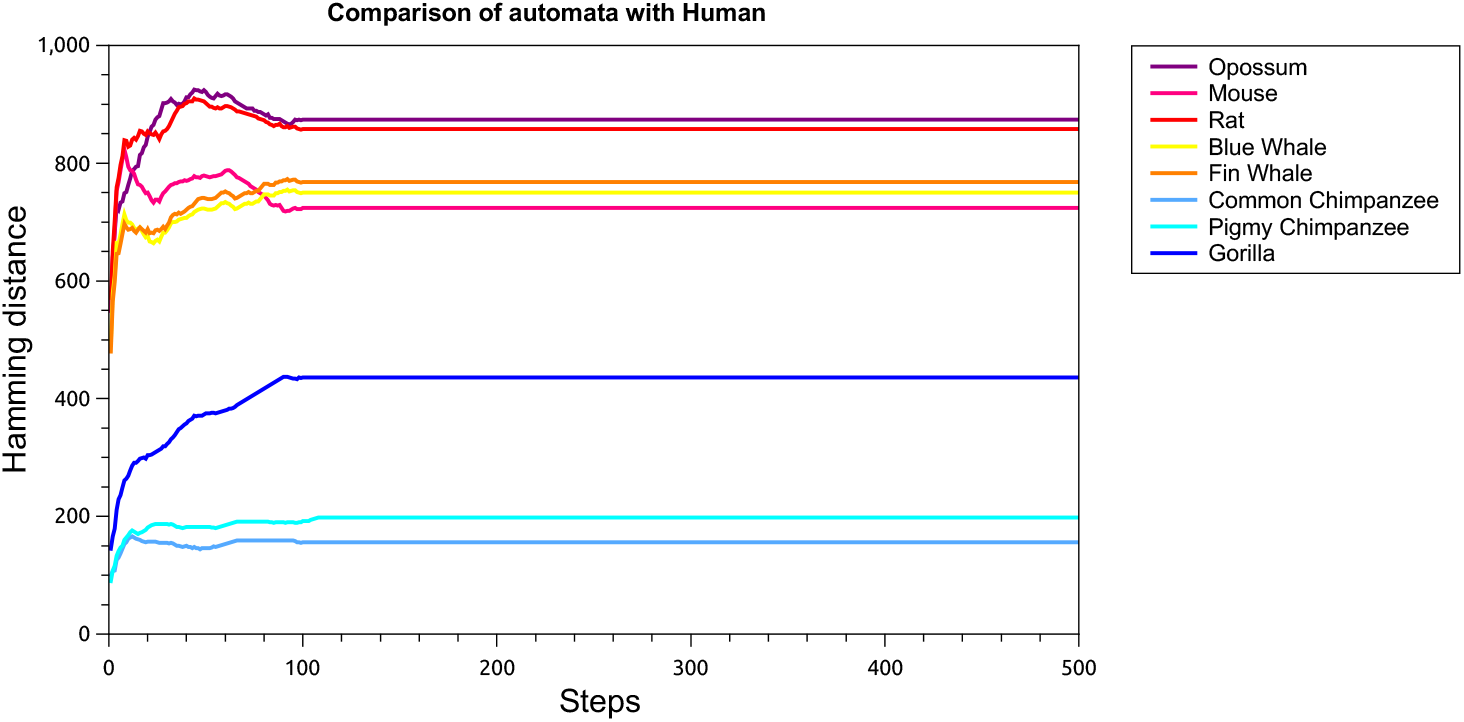
Hamming distance between the cellular automata image for some different mammalian species and Human, as a function of the number of steps.

We then build distance matrices from the SHD and the p-distance, and obtain corresponding dendrograms using the average method from R studio hierarchical grouping [38, 39], and shown in Fig. 5. Both dendrograms are identical, with the exception of a small difference of the closest relative of human. Nevertheless both methods correctly group families: Hominidae, Balaenopteridae, Muridae, and Didelphidae. Other methods such as [35] yield dendrograms identical to the one obtained from our method. The cophenetic correlation coefficient [40] between the two dendrogram in Fig. 5 is 0.9940, indicating a very close similarity.

**Fig 5.**
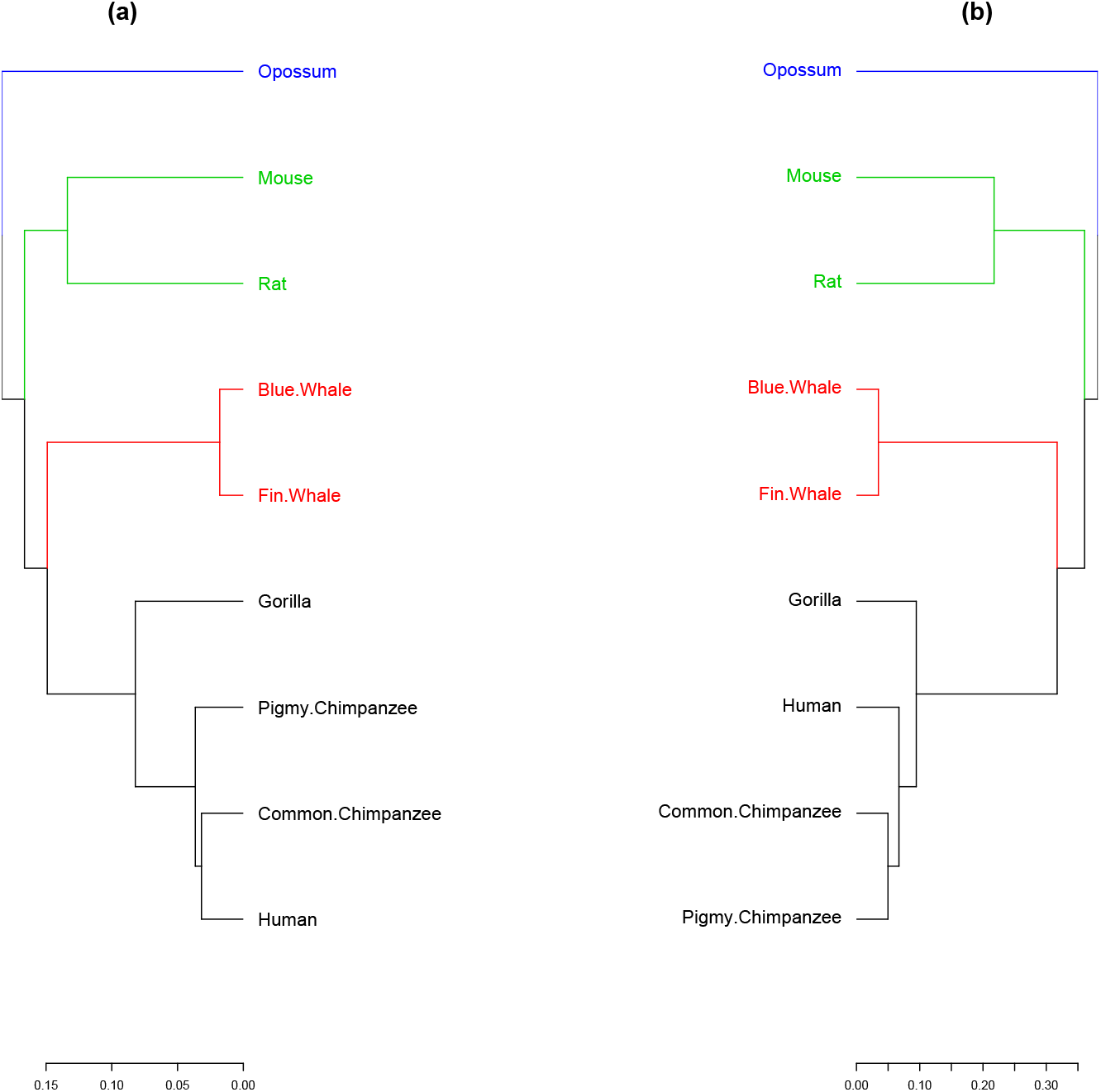
Dendrograms from the ND 5 protein from (a) SHD and (b) p-distance. The four families are well grouped in both dendrograms: Didelphidae (blue), Muridae (green), Balaenopteridae (red), and Hominidae (black).

### NADH Dehydrogenase 6 (ND 6)

The NADH Dehydrogenase 6 (ND 6) protein is a sub-unit of the NADH dehydrogenase (ubiquinone) enzyme, located in the mitochondrial inner membrane. Mutations or errors in their sequences can cause Leigh’s disease and spinal muscular atrophy [41]. Protein sequences were obtained from NCBI [6], and their identifications are given in S2 Table.

The aligned sequences have 176 positions, and thence the initial condition for this protein has 8 *×* 176 = 1408 cells. We follow the same procedure as for the previous case: the SHD between each pair among the following species: Human, Gorilla, Common Chimpanzee, Gray Seal, Harbor Seal, Rat, Mouse, and Wallaroo. Dendrograms are then obtained from the distance matrices using the SHD and the p-distance, and shown in Fig. 6. The two dendrograms are identical and both methods group families correctly: Macropodidae, Muridae, Phocidae, and Hominidae. We note that other methods as alignment-free similarity analysis [35] cannot separate the Macropodidae family from the Muridae. The cophenetic correlation coefficient between the two dendrograms in Fig. 6 is 0.9797, indicating again a very close similarity.

**Fig 6.**
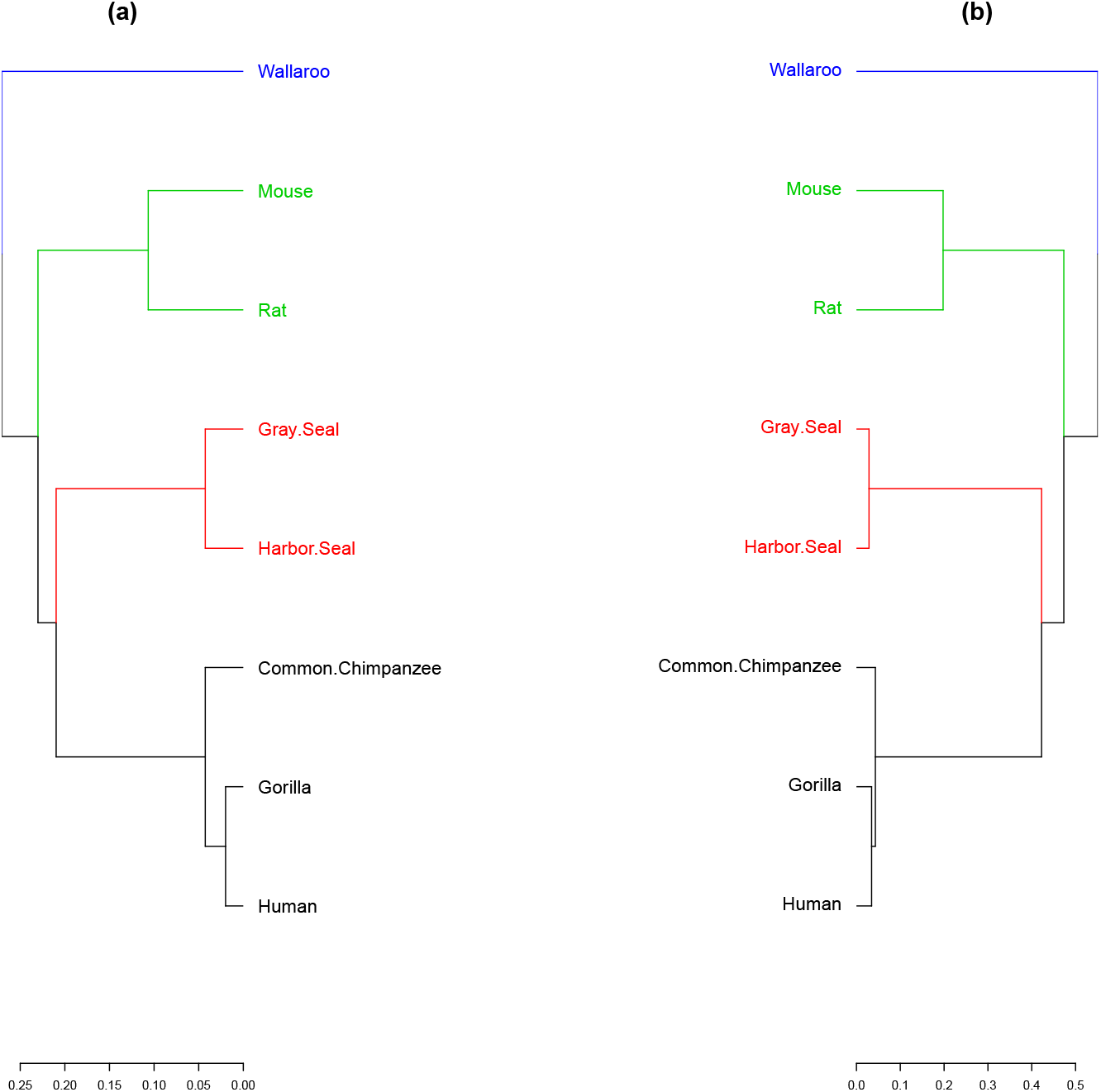
Dendrograms from the ND 6 protein obtained from distance matrices using (a) SHD and (b) p-distance, with the indication of family groupings: Macropodidae (blue), Muridae (green), Phocidae (red), and Hominidae(black).

### Transferrin

Transferrin is an iron-binding protein keeping iron at a low concentration in biological fluids. Serum transferrin (TF) is present in mammals, amphibians, and fish [42], and plays an essential role in fighting bacterial infections in fish [43]. Blood iron overload is a rare condition that characterizes hereditary atransferrinemia [44]. We consider a set of 24 transferrin protein sequences across Mammalia, Amphibian, and Actinopterygii species from the NCBI database [6] and their identifications are given in S3 Table. The aligned sequences have 750 positions. Each one is then encoded into a binary code of size 8 *×* 750 = 6000. The dendrogram obtained from the SHD and p-distance are shown in Fig. 7. Our approach correctly classifies all species into their respective groups: Mammalia, Amphibian and Actinopterygii and separately grouping mammals’ serum transferrin (TF) and lactotransferrin (LF). It also group correctly species from the genus Salmo (Brown Trout, Atlantic Salmon), Salvelinus (Japanese Char, Brook Trout, Lake Trout), and Oncorhynchus (Amago Salmon, Sockeye Salmon, Rainbow Trout, Coho Salmon, Chinook Salmon) in the Actinopterygii class. Only Amago Salmon (TF) and Sockeye Salmon (TF) were grouped differently, but the same problem has already been reported in previous works [43]. The cophenetic correlation coefficient between the two dendrograms is 0.9671, again indicating a very good similarity between the clusters.

**Fig 7.**
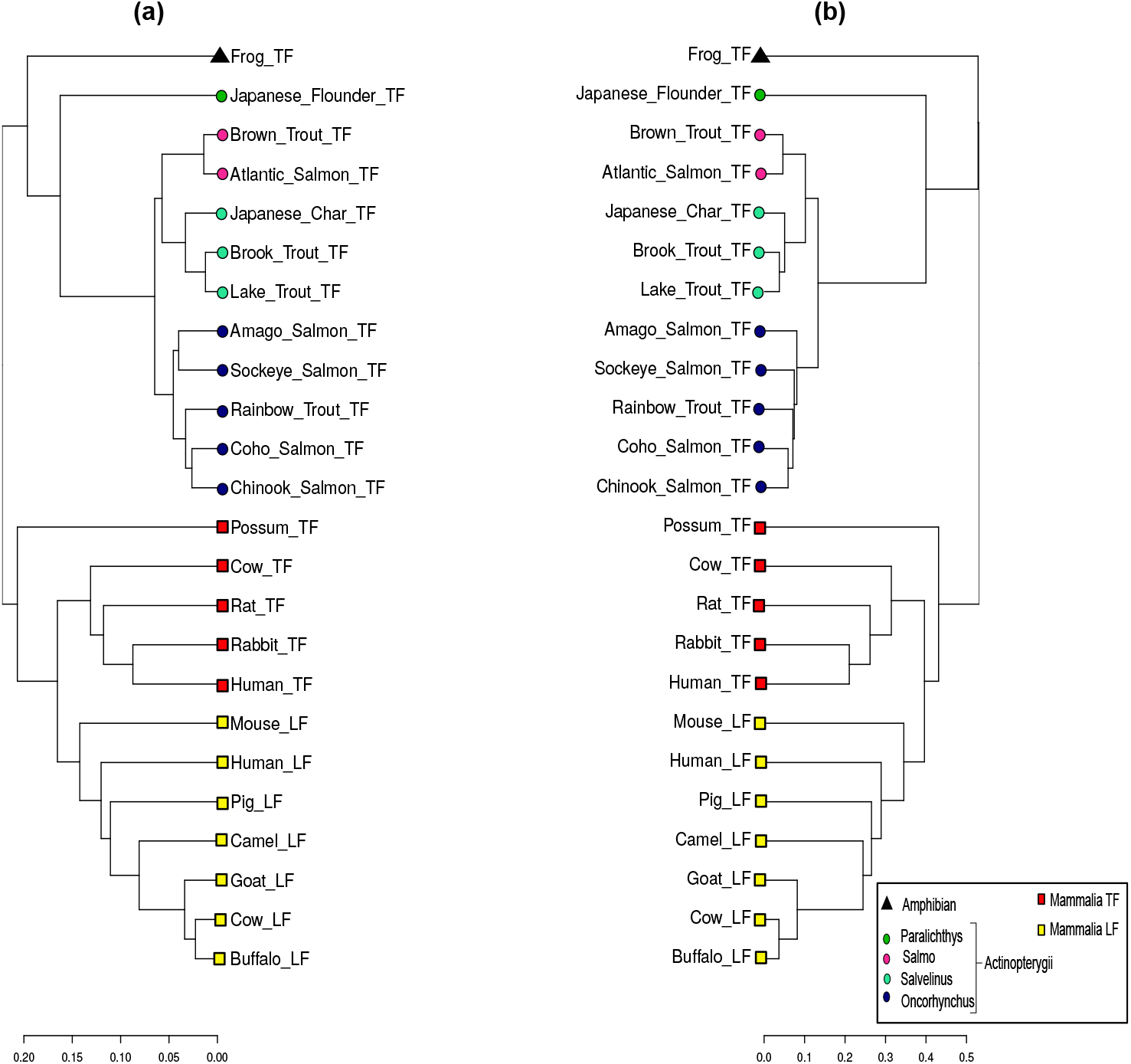
Transferrin protein dendrograms from (a) SHD and (b) p-distance distance matrices.

### Beta-Globin

Hemoglobin comprises two chain pairs *α* and *β*, which have distinct chains of amino acids, two dimers of *α − β* form hemoglobin. Its principal function is to carry oxygen from blood to tissues. Mutations in the beta-globin chain can cause sickle cell anemia [45]. We consider here 50 beta-globin sequences from different species taken from the NCBI database [6], and their identifications are given in S4 Table. The aligned sequences have 148 positions. The initial condition is then coded in 8 *×* 148 = 1184 cell digits. The corresponding distance matrices for SHD and p-distance then have 50 *×* 50 = 2500 entries, and the corresponding dendrograms are shown in Fig. 8.

**Fig 8.**
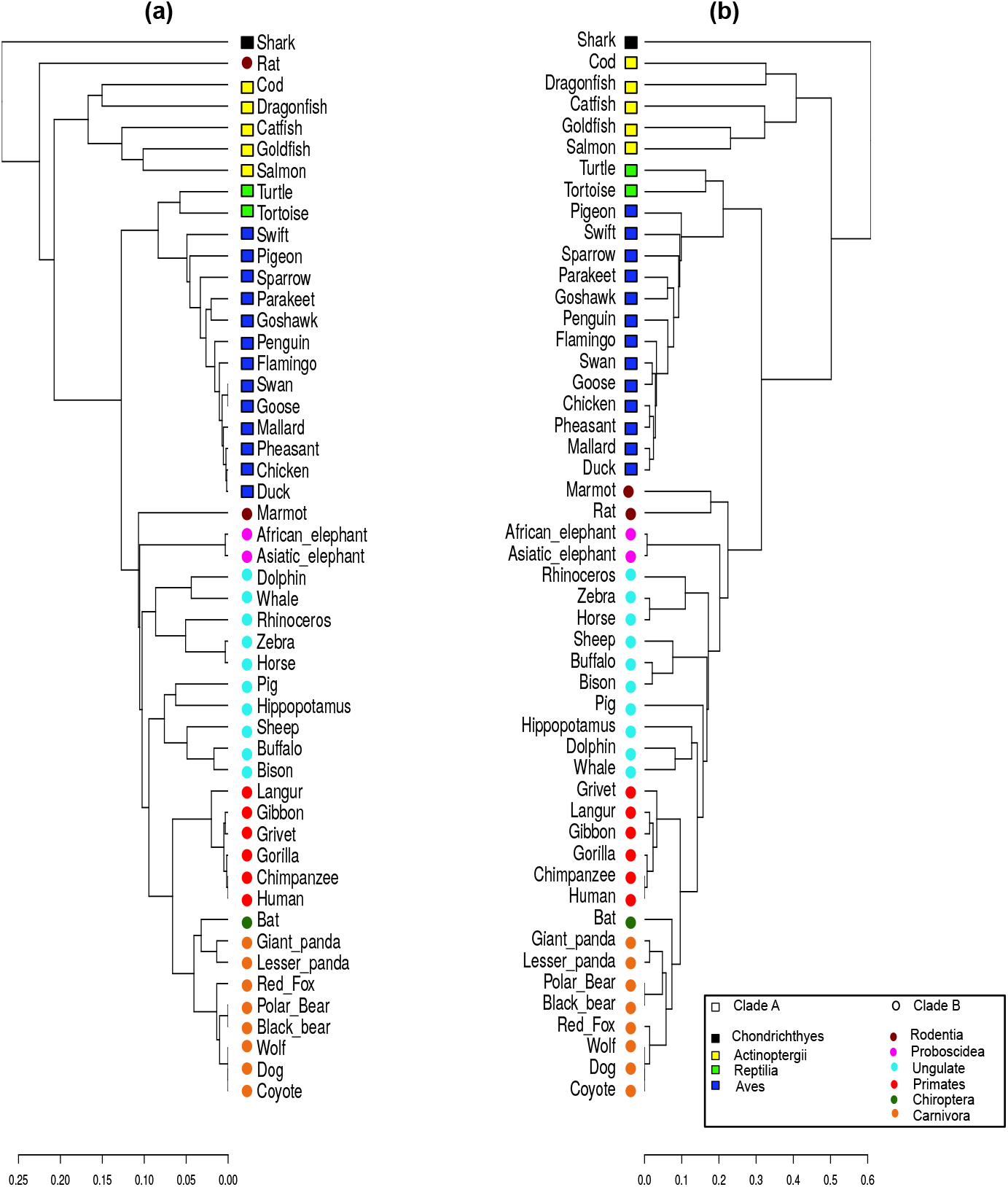
Beta-globin protein dendrograms from SHD (a) and p-distance (b) values. Clade A is represented by squares and Clade B by circles. Different animal groups are represented by colors.

Our approach yields a consistent classification of the identified clusters. At variance the results in [13] and the dendrogram obtained using the p-distance, that separates mammals from non-mammals, our approach failed in this point, with rat classified in Clade A. This same inconsistency was also observed in a previous work [35]. Other divergences of the method involve some more recent families but keep the tree similar to the one obtained using the p-distance. These discrepancies observed for Beta-Globin may be due to the fact that we considered a significant and more diverse number of species, and is reflected in the value of 0.8790 for the cophenetic correlation coefficient between the two dendrograms, clearly below the other proteins considered here, but still acceptable.

## Conclusion

We discussed and showed that Cellular automata are a tool for visual comparing of protein sequences and for determining their similarity. We expanded the use of this tool by introducing the use of the Hamming distance from information theory, in order to compare the cellular automata images obtained. Our approach allows to determine phylogenetic relations among species with a good accuracy if one considers that one protein was used in each of the dendrogram presented, but nevertheless has some limitations. We applied it to lysozyme protein sequences (not shown here), with inconclusive results, with the possible explanation that the sequences fir this cases not homologous but are the result of convergent evolution. In this case, the resulting dendrograms from both our method and by using p-distance cannot approximate species with recent common ancestors.

The main advantage of our method compared to using the p-distance to build the distance matrix is that it codes each amino acid according to its structure, such that similar amino acids have closer codings, such that the distance measure we use will give different weights for different mutations, as observed in [28], which used a code based on the hydrophobicity of amino acids such that the sequences that underwent mutations with a change in hydrophobicity had a greater distance and also a different weight for each type of mutation.

Other approaches use textures from images [21] to compare cellular automata. Ours requires a low computational cost and no processing methods or image textures, with an efficient protein comparison. As a first work we used an evolution rule previously proposed in the literature, but in forthcoming research we will consider other possibilities, and are currently investigating the possibility of coding proteins using the hydrophobicity scale proposed by Moret and Zebende [46].

## Supporting information

All sequence data were taken from the NCBI genome database https://www.ncbi.nlm.nih.gov/protein. Below are the identifications of each species and protein used.

**S1 Table.** Identification of ND5 protein sequences at NCBI.

| Sequence name | Species | ID (NCBI) |
| --- | --- | --- |
| Human | Homo sapiens | AP 000649.1 |
| Gorilla | Gorilla gorilla | NP 008222.1 |
| Common chimpanzee | Pan troglodytes | NP 008196.1 |
| Pigmy chimpanzee | Pan paniscus | NP 008209.1 |
| Fin whale | Balenoptera physalus | NP 006899.1 |
| Blue whale | Balenoptera musculus | NP 007066.1 |
| Rat | Rattus norvegicus | AP 004902.1 |
| Mouse | Mus musculus | NP 904338.1 |
| Opossum | Didelphis virginiana | NP 007105.1 |

**S2 Table.** Identification of ND6 protein sequences at NCBI.

| Sequence name | Species | ID (NCBI) |
| --- | --- | --- |
| Human | Homo sapiens | YP 003024037.1 |
| Gorilla | Gorilla gorilla | NP 008223.1 |
| Common chimpanzee | Pan troglodytes | NP 008197.1 |
| Harbor seal | Phoca vitulina | NP 006939.1 |
| Gray seal | Halichoerus grypus | NP 007080.1 |
| Rat | Rattus norvegicus | AP 004903.1 |
| Mouse | Mus musculus | NP 904339.1 |
| Wallaroo | Osphranter robustus | NP 007405.1 |

**S3 Table.** Identification of Transferrin protein sequences at NCBI.

| Sequence name | Species | ID (NCBI) |
| --- | --- | --- |
| Human TF | Homo sapiens | S95936.1 |
| Rabbit TF | Oryctolagus cuniculus | P19134.4 |
| Rat TF | Rattus norvegicus | D38380.1 |
| Cow TF | Bos Taurus | U02564.1 |
| Buffalo LF | Bubalus bubalis | AJ005203.1 |
| Cow LF | Bos Taurus | X57084.1 |
| Goat LF | Capra hircus | X78902.1 |
| Camel LF | Camelus dromedarius | AJ131674.1 |
| Pig LF | Sus scrofa | AAA31102.1 |
| Human LF | Homo sapiens | NM 002343.6 |
| Mouse LF | Mus musculus | NM 008522.3 |
| Possum TF | Trichosurus vulpecula | AF092510.1 |
| Frog TF | Xenopus laevis | X54530.1 |
| Japanese flounder TF | Paralichthys olivaceus | D88801.1 |
| Atlantic salmon TF | Salmo salar | L20313.1 |
| Brown trout TF | Salmo trutta | D89091.1 |
| Lake trout TF | Salvelinus namaycush | D89090.1 |
| Brook trout TF | Salvelinus fontinalis | D89089.1 |
| Japanese char TF | Salvelinus leucomaenis pluvius | D89088.1 |
| Chinook salmon TF | Oncorhynchus tshawytscha | AH008271.2 |
| Coho salmon TF | Oncorhynchus kisutch | D89084.1 |
| Sockeye salmon TF | Oncorhynchus nerka | D89085.1 |
| Rainbow trout TF | Oncorhynchus mykiss | D89083.1 |
| Amago salmon TF | Oncorhynchus masou | D89086.2 |

**S4 Table.** Identification of Beta-Globin protein sequences at NCBI.

| Sequence name | Species | ID (NCBI) |
| --- | --- | --- |
| Human | Homo sapiens | AAA16334.1 |
| Pigeon | Columba Livia | P11342.1 |
| Goshawk | Accipiter gentilis | P08851.1 |
| Black Bear | Ursus thibetanus | P68012.1 |
| Lesser Panda | Ailurus fulgens | P18982.1 |
| Asiatic Elephant | Elephas maximus | P02084.1 |
| Giant Panda | Ailuropoda melanoleuca | P18983.2 |
| African Elephant | Loxodonta africana | P02085.1 |
| Sheep | Ovis aries | P02075.2 |
| Tortoise | Chelonoidis niger | P83123.3 |
| Duck | Anas platyrhynchos | P02114.2 |
| Grivet | Chlorocebus aethiops | P02028.1 |
| Mallard | Anas platyrhynchos platyrhynchos | P02115.1 |
| Gorilla | Gorilla gorilla gorilla | P02024.2 |
| Goose | Anser anser anser | P02117.1 |
| Shark | Heterodontus portusjacksoni | P02143.1 |
| Rat | *Rattus norvegicus* | CAA33114.1 |
| Hippopotamus | *Hippopotamus amphibius* | P19016.1 |
| Penguin | *Aptenodytes forsteri* | P80216.1 |
| Horse | *Equus caballus* | P02062.1 |
| Swift | *Apus apus* | P15165.1 |
| Gibbon | *Hylobates lar* | P02025.1 |
| Coyote | *Canis latrans* | P60525.1 |
| Whale | *Balaenoptera acutorostrata* | P18984.1 |
| Catfish | *Silurus asotus* | O13163.2 |
| Bat | *Macroderma gigas* | P24660.1 |
| Bison | *Bison bonasus* | P09422.1 |
| Red Fox | *Vulpes vulpes* | P21201.1 |
| Swan | *Cygnus olor* | P68945.1 |
| Marmot | *Marmota marmota* | P08853.1 |
| Buffalo | *Bubalus bubalis* | P67820.1 |
| Salmon | *Salmo salar* | Q91473.3 |
| Dog | *Canis lupus familiaris* | P60524.1 |
| Sparrow | *Passer montanus* | P07406.1 |
| Chimpanzee | *Pan troglodytes* | P68873.2 |
| Pheasant | *Phasianus colchicus colchicus* | P02113.1 |
| Dolphin | *Tursiops truncatus* | P18990.1 |
| Flamingo | *Phoenicopterus ruber* | P02121.1 |
| Goldfish | *Carassius auratus* | P02140.1 |
| Pig | *Sus scrofa* | P02067.3 |
| Polar bear | *Ursus maritimus* | P68011.1 |
| Dragonfish | *Cygnodraco mawsoni* | ADD73488.1 |
| Rhinoceros | *Rhinoceros unicornis* | P09907.1 |
| Parakeet | *Psittacula krameri* | P21668.1 |
| Chicken | *Gallus gallus* | P02112.2 |
| Zebra | *Equus zebra* | P67824.1 |
| Wolf | *Chrysocyon brachyurus* | P60526.1 |
| Cod | *Gadus morhua* | O13077.2 |
| Turtle | *Chrysemys picta bellii* | P13274.1 |
| Langur | *Semnopithecus entellus* | P02032.1 |

## Notes

### Competing Interest Statement

The authors have declared no competing interest.

